# Testing the Gene Expression Classification of the EMT Spectrum

**DOI:** 10.1101/452508

**Authors:** Dongya Jia, Jason T. George, Satyendra C. Tripathi, Deepali L. Kundnani, Mingyang Lu, Samir M. Hanash, José N. Onuchic, Mohit Kumar Jolly, Herbert Levine

## Abstract

The epithelial-mesenchymal transition (EMT) plays a central role in cancer metastasis and drug resistance – two persistent clinical challenges. Epithelial cells can undergo a partial or full EMT, attaining either a hybrid epithelial/mesenchymal (E/M) or mesenchymal phenotype, respectively. Recent studies have emphasized that hybrid E/M cells may be more aggressive than their mesenchymal counterparts. However, mechanisms driving hybrid E/M phenotypes remain largely elusive. Here, to better characterize the hybrid E/M phenotype(s) and tumor aggressiveness, we integrate two computational methods – (a) RACIPE – to identify the robust gene expression patterns emerging from the dynamics of a given gene regulatory network, and (b) EMT scoring metric - to calculate the probability that a given gene expression profile displays a hybrid E/M phenotype. We apply the EMT scoring metric to RACIPE-generated gene expression data generated from a core EMT regulatory network and classify the gene expression profiles into relevant categories (epithelial, hybrid E/M, mesenchymal). This categorization is broadly consistent with hierarchical clustering readouts of RACIPE-generated gene expression data. We show that the EMT scoring metric can be used to distinguish between samples composed of exclusively hybrid E/M cells and those containing mixtures of epithelial and mesenchymal subpopulations using the RACIPE-generated gene expression data.

## Introduction

The epithelial-mesenchymal transition (EMT) is a trans-differentiation developmental program by which epithelial (E) cells weaken their adhesive bonds with neighbors as well as apico-basal polarity, while concomitantly gaining mesenchymal (M) traits of migration and/or invasion [1,2]. Cancer cells undergoing EMT typically have enhanced metastatic abilities [2,3], elevated resistance against chemotherapy [4,5] and can evade the immune system [6], thus accelerating malignancy. In the developmental context of mammary morphogenesis and the physiological context of wound healing, cells rarely undergo a complete EMT. Instead, they exhibit a transient hybrid epithelial/mesenchymal (E/M) phenotype that shows a mixture of traits - epithelial traits of cell-cell adhesion, and mesenchymal traits of migration [3]. Thus, the hybrid E/M phenotype facilitates collective cell migration, instead of individual cell migration - a characteristic of cells having undergone a complete EMT. This concept has recently gained momentum in the context of cancer metastasis, wherein it has been observed that malignant cells can maintain a hybrid E/M phenotype and migrate collectively, eventually appearing in the blood as clusters of circulating tumor cells (CTCs) [7–10]. Moreover, these CTC clusters, often five to eight cells in size, contribute much more than their proportional share to metastases [11], thus behaving as ‘chief instigators’ of metastasis. Cells in these clusters often display a hybrid E/M phenotype [9,10,12], and can resist cell death in circulation by evading immune attacks and providing survival signals to one another through their cell-cell contacts [12]. Therefore, hybrid E/M cells may occupy the ‘metastatic sweet spot’ [13,14].

The hybrid E/M phenotype has been tacitly assumed to be ‘metastable’ or transient [15,16], but recent computational studies suggest that the hybrid E/M phenotype can be a stable cell phenotype, particularly in the presence of ‘phenotypic stability factors’ (PSFs) such as the transcription factors GRHL2, OVOL2, NUMB and ∆Np63α [10,17–22]. Resonating with predictions of these computational models, non-small cell lung cancer (NSCLC) cells H1975 have been shown to stably (i.e. over multiple passages) co-express epithelial and mesenchymal markers - E-cadherin and vimentin - at a single-cell level *in vitro* [19]. Consistent with predictions from mathematical models, knockdown of GRHL2, OVOL2 or NUMB in H1975 cells results in a complete EMT [18,19,22]. Overall, these studies have supported the relevance of a stable hybrid E/M phenotype and PSFs along with their implications for both mammary morphogenesis and cancer metastasis [12].

EMT is orchestrated by complex gene regulatory networks (GRNs) that involve microRNAs (miRs), transcription factors (TFs), alternative splicing mechanisms, and epigenetic modifications [23,24]. Particularly, members of the miR-200 and the miR-34 family can maintain or induce epithelial features, and TFs such as ZEB1/2, SNAI1/2 and FOXC2 can trigger the transition toward a mesenchymal phenotype.

Moreover, these regulators exhibit extensive crosstalk, forming multiple feedback loops [23]. Mathematical modeling approaches have been used to analyze the dynamics of EMT regulatory networks [17,25–31]. However, most of these studies consider interactions among only a small number of genes regulating EMT, because a quantitative simulation of the GRN dynamics is difficult owing to insufficient knowledge of kinetic parameter values, particularly when the network size is large.

Recently, we developed a computational method, <u>ra</u>ndom <u>ci</u>rcuit <u>pe</u>rturbation (RACIPE), to interrogate the robust dynamical features of GRNs without the need for precise parameter values [32,33]. RACIPE generates an unbiased ensemble of mathematical models based on the topological information (i.e. genes and interactions) of a GRN. Each model is characterized by a unique set of parameters that are randomly sampled within ranges of values that are biologically relevant, and then numerically solved. The stable state solutions of the ensemble models are collected and analyzed, through which the most significant gene expression patterns associated with a specific GRN are extracted. In this framework, variability among different sets of parameters generated by RACIPE captures both the heterogeneity among individual cells and the effects of different input signals to the GRN. Thus, the RACIPE-generated stable state solutions can, in principle, represent the gene expression patterns of diverse individual cell phenotypes in multiple contexts. Here, we apply RACIPE to EMT regulatory networks to generate *in silico* gene expression data of different EMT phenotypes.

Separately, the lack of a rigorous quantitative definition of a hybrid E/M phenotype motivated us to develop a statistical learning method, the EMT scoring metric [34]. This approach assigns a probability of membership in the hybrid E/M category, given the gene expression data of a sample. It also assigns each sample a corresponding EMT score on a scale from 0 (fully epithelial) to 2 (fully mesenchymal) with an intermediate score of 1 representing maximally hybrid E/M signatures. This approach, when applied to the NCI-60 human tumor cell line gene expression dataset as the training set, identified claudin-7 (CLDN7) and the ratio of vimentin to E-cadherin (VIM/CDH1) as the best-fit pair of predictors to categorize the EMT phenotypes. The algorithm has been accurate in characterizing the EMT status of multiple cancer cell lines and has elucidated a subtype-specific association between the EMT status and patient survival.

Here, we use RACIPE to test the ability of our EMT scoring metric to quantify the EMT phenotypes. We first apply RACIPE on a proposed core EMT regulatory network, containing 26 genes and 100 regulatory links (**Fig. 1a**) and use the hierarchical clustering analysis (HCA) to classify the RACIPE-generated gene expression data into four main clusters corresponding to one epithelial, one mesenchymal, and two hybrid E/M (E/M I and II) phenotypes. We show that the EMT scoring metric can accurately recapitulate the EMT status of RACIPE-generated hybrid E/M samples. We next construct *in silico* mixtures of RACIPE-generated E, M and E/M II samples in various ratios. We present a proof-of-principle for distinguishing samples containing individual hybrid E/M cells from those containing mixtures of E and M cells by applying an extension of our EMT scoring metric. We demonstrate that the EMT metric approach can accurately identify the proportions of the different subpopulations for all the various combinations of mixtures studied here. This new method is enabled by having samples generated from RACIPE projected on to the EMT scoring metric space. Finally we validate the proposed method comparing its prediction to data from multiple non-small cell lung cancer (NSCLC) cell lines.

## Materials and Methods

### Generation of EMT gene expression data by RACIPE

The EMT regulatory network (**Fig. 1a**) we consider here contains 26 gene products and 100 regulatory links, the majority of which are transcriptional in nature. There are also microRNA-mediated translational regulation. All links in the network considered here are modeled by shifted Hill functions [25]. RACIPE takes the topological information of the EMT regulatory network as the input and generates multiple mathematical models.

The general rate equation representing the temporal dynamics for a single gene is of the form:

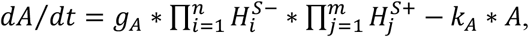

where *A* represents the expression level of gene A (the level of mRNA A), *g_A_* and *k_A_* represent the innate production and degradation rates of mRNA A, 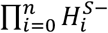 and 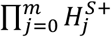 represent the inhibitory regulation and excitatory regulation of gene A by other genes, respectively; the number of inhibitory links is *n* and the number of excitatory links is *m. H^S−^* and *H^S+^* are negative and positive shifted Hill functions of the form:

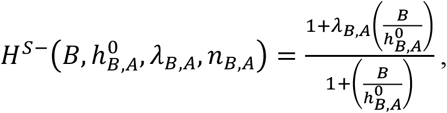

and

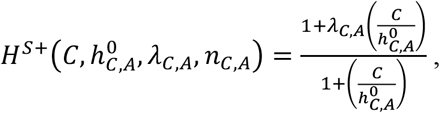

where *λ_B,A_* < 1 is the fold-change representing the inhibition of gene A expression by the protein encoded by gene B, *λ_C,A_* > 1 is the fold-change representing the activation of gene A by the protein encoded by gene C, *B* represents the levels of mRNA B, and *C* represents the levels of mRNA C. 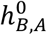 and 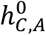 are thresholds in the shifted Hill function.

The temporal dynamics of all 26 nodes of the EMT regulatory network are characterized by the above functional form. In the RACIPE framework, mathematical model refers to a specific choice of parameters, and many models are created. We typically use 10,000 to 100,000 models. For each model, the parameters are randomly sampled within biologically reasonable ranges. The sampled parameters satisfy the ‘half-functional’ rule [32] so that each regulatory link has an equal chance (~50%) to be active or inactive. For generating each mathematical model, 200 initial conditions are used to numerically integrate the dynamical equations using Euler’s Method and we keep all simulations for which the system reaches steady state on the chosen integration interval. Depending on the specific parameters, a single model may give rise to one or more stable steady state solutions dependent upon the model’s initial conditions. The stable steady state solutions from all models are analyzed via the EMT scoring metric.

### EMT scoring metric

In this approach, statistical models are created to resolve training samples into their *a priori* known categories. EMT scoring is performed by applying the best-fit model obtained from an iterative statistical procedure developed previously to relevant gene expression samples [34]. This best fit model was found to maximally resolve the NCI-60 cell line gene expression dataset (GSE5846) based on E, M, or hybrid E/M phenotype using ordinal multinomial logistic regression. The (log2-normalized) input predictors include the ratio of vimentin and E-cadherin (VIM/CDH1) and claudin 7 (CLDN7). Model output for a given sample, *S*, is given by

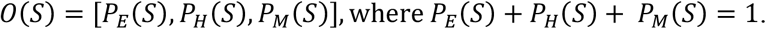

This ordered triplet is interpreted as the probability that sample *S* falls into either of the following three categories, epithelial (E), hybrid E/M (H) and mesenchymal (M), *P_E_*(*S*), *P_H_*(*S*), *P_M_*(*S*). Ordinal regression places state H intermediary to E and M, and the EMT metric, *μ*, assigns a numeric value on the interval [0, 2] with the ordering *E < H < M*. In particular,

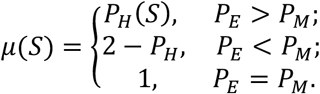

The metric represents epithelial samples in the range 0 ≤ *μ* < 0.5, hybrid for 0.5 ≤ *μ ≤* 1.5, and mesenchymal for 1.5 < *μ ≤2*.

### Generating the mixtures of E, M and hybrid E/M II samples

We generate mixtures of E and M samples with three replicates of five different ratios of E to M: [50%, 50%], [20%, 80%], [80%, 20%], [40%, 60%] and [60%, 40%] (n=20,000 per ratio per replicate). For example, 10^4^ E samples are randomly sampled from RACIPE-generated E samples and 10^4^ M samples are randomly sampled from RACIPE-generated M samples and their mixture has a proportion of E and M -[50%, 50%]. The mean gene expression of the mixture is used to estimate the proportions of E and M by the EMT metric. In a similar way, mixtures of E and hybrid E/M II samples and those of M and hybrid E/M II samples with three replicates of five different proportions - [50%, 50%], [60%, 40%], [40%, 60%], [20%, 80%] and [80%, 20%] - are generated respectively (n=15,000 per ratio per replicate). Mixtures of E, M and hybrid E/M II samples with three replicates of five different proportions - [25%, 25%, 50%], [30%, 30%, 40%], [20%, 20%, 60%], [10%, 10%, 80%] and [40%, 40%, 20%] - are generated respectively (n=15,000 per ratio per replicate).

### Predictions of the mixture proportions

Samples are depicted by their location in the two-dimensional EMT predictor space. This space is spanned by VIM/CDH1 and CLDN7 expression levels partitioned into three regions based on which one of {P_E_(S), P_H_(S), P_M_(S)} is maximal. For example, when investigating mixtures of E and M populations, predictions of the average proportion of E and M cells are obtained by projecting the coordinates of a sample (x_S_, y_S_) to the curve adjoining the location of the mean E signature at (x_E_,y_E_), and mean M signature at (x_M_, y_M_) (x and y correspond to mean CLDN7 and (mean VIM)/(mean CDH1) levels, respectively). The proportion of E and M for sample (x_S_, y_S_) is estimated by finding the least-squares projection of the sample onto the curve spanned by the convex combination of E and M signatures. The mixture proportions of E and hybrid E/M II populations and the mixture proportions of M and hybrid EM II populations are predicted in an identical way. Mixtures of equal parts E and M with variable proportions of hybrid E/M II samples were analyzed in a similar manner, instead using convex signatures of E/M II and the midpoint on the convex curve adjoining E and M.

### Cell culture and immunofluorescence

NSCLC cells were cultured in RPMI 1640 medium containing 10% fetal bovine serum and 1% penicillin/streptomycin cocktail (Thermo Fisher Scientific, Waltham, MA). For immunofluorescence, cells were fixed in 4% paraformaldehyde, permeabilized in 0.2% Triton X-100, and then stained with anti-CDH1 (1:100; Abcam) and anti-vimentin (1:100; Cell Signaling Technology). The primary antibodies were then detected with Alexa conjugated secondary antibodies (Life technologies). Nuclei were visualized by co-staining with DAPI.

## Results

### Construction of the regulatory network of EMT

Based on an extensive literature search, we construct a core gene regulatory network of EMT including nine microRNAs (miR-141, miR-200a,b,c, miR-34a, miR-101, miR-30c, miR-9, miR-205), ten EMT-inducing transcription factors (EMT-TFs) (ZEB1, ZEB2, FOXC2, SNAI1, SNAI2, TWIST1, TWIST2, GSC, KLF8, TCF3), one EMT-inducing signal (TGF-β), three PSFs (OVOL2, GRHL2 and ∆Np63α), CLDN7 and the canonical epithelial and mesenchymal markers CDH1 and VIM respectively. The network of EMT presented here is an extended version of the one used in our previous study [32]. Specifically, three proposed PSFs, OVOL2, GRHL2 and ∆Np63α, together with CLDN7 are included due to their identified roles in epithelial-mesenchymal plasticity (See **SI Section 1** and **Table S1** for more information).

### RACIPE identifies the gene expression patterns of multiple EMT phenotypes

We apply RACIPE to the core gene regulatory network above (**Fig. 1a**) and collect the gene expression data from 10,000 different sets of parameters (10,000 distinct RACIPE models). Hierarchical clustering analysis, used to analyze the gene expression patterns of RACIPE-generated data, reveals four large clusters (**Fig. 1b**). The first cluster is characterized by high expression of epithelial-associated genes or microRNAs, such as miR-200, miR-34, CLDN7 and CDH1, and low expression of mesenchymal-associated genes, such as ZEB1, FOXC2, SNAIL and VIM, thus identified as an epithelial signature. The second cluster shows low expression of epithelial-associated genes or microRNAs and high expression of mesenchymal-associated genes and therefore represents a mesenchymal signature. The remaining two clusters have co-expression of epithelial-associated and mesenchymal-associated genes to varying extents, and so these two clusters are characterized as hybrid E/M I and II signatures. Notably, the PSFs GRHL2, OVOL2 and ∆Np63α are expressed at high levels in hybrid E/M I samples, but not in hybrid E/M II samples. In contrast, miR-101 and GSC are expressed at high levels in hybrid II samples while not in hybrid E/M I samples. The distinct gene expression profiles of hybrid E/M I and hybrid E/M II shown here emphasize the heterogeneity of one or more hybrid E/M phenotype(s) that cells may attain during EMT [35–37].

**Figure 1.**
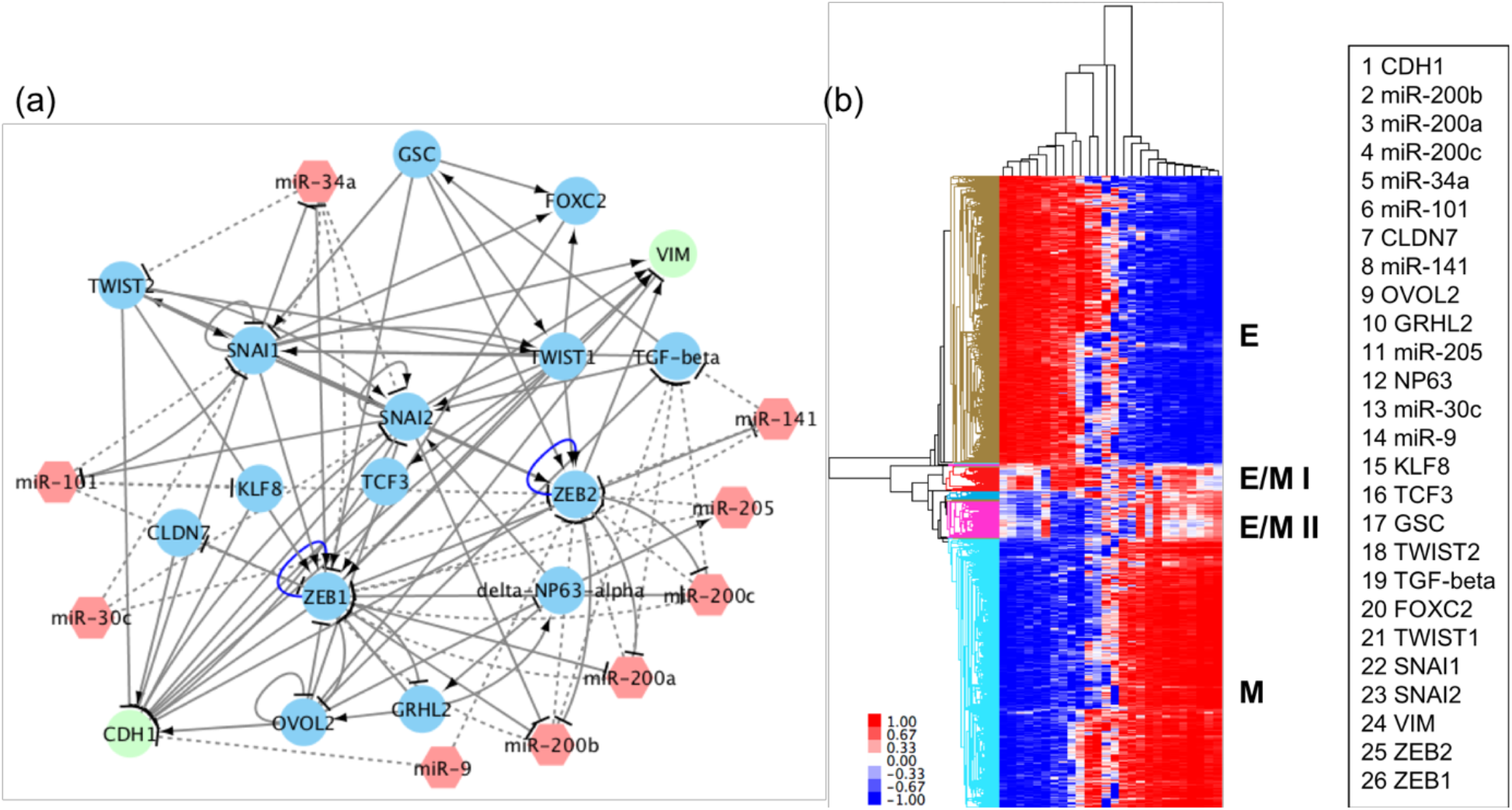
**RACIPE identified robust gene expression patterns of EMT**. (a) A proposed core EMT regulatory network. The EMT regulatory network contains 9 microRNAs (red hexagons) and 17 non-microRNAs (circles). Solid arrows represent excitatory regulation, mostly due to transcriptional activators. Solid bar-headed arrows represent inhibitory regulation, mostly due to transcriptional inhibitors. Dotted bar-headed arrows represent microRNA-mediated regulations. (b) Hierarchical clustering analysis of the RACIPE-generated gene expression data from 10,000 RACIPE models. Each row represents one sample, i.e., one RACIPE-generated stable steady state, and each column represents one gene. The names of genes corresponding to each column from left to right are listed.

### The EMT scoring metric captures the hybrid E/M I and II signatures generated by RACIPE

The heterogeneity of RACIPE-generated aggregate output may be appreciated by projection of these samples onto the two-dimensional EMT metric predictor space (**Fig. 2a**). This space is divided into three colored areas: blue (mesenchymal), dark green (hybrid E/M), and red (epithelial). Samples belong to a particular category with high probability if they localize far from the boundary between two regions. We observe high variability within each phenotypic cluster when projecting RACIPE-generated data on this predictor space, which is unsurprising given that solutions are generated from an ensemble of randomized parameters. Thus, we focus our attention to the average (mean) values of individual genes for all the samples in a given category, referred to as the mean signatures. The mean signatures of hybrid E/M I and E/M II samples are appropriately binned via the EMT metric — the probability that hybrid E/M I is categorized as hybrid E/M is 0.76 with intermediate EMT score of *μ* = 0.76. and the probability of hybrid categorization for hybrid E/M II is 0.88 with intermediate EMT score of *μ* = 1.12. The probability of hybrid categorization may be visually appreciated in **Fig. 2b**, wherein samples with larger probabilities of belonging to hybrid E/M localize inside the green region further from the interface between other groups. Our findings highlight the consistency between RACIPE-generated gene expression samples for hybrid E/M phenotypes and their expected location on the EMT spectrum as generated by our predictive metric. The mean signature of mesenchymal samples is also captured by the EMT metric with a 0.71 probability of being binned as mesenchymal, and a high EMT score of *μ* = 1.71. (**Fig. 2b**). However, the mean signature of epithelial samples has a slightly higher probability of being binned as hybrid E/M (*P_H_* = 0.59) as compared to epithelial (*P_E_* = 0.41) with an EMT score of *μ* = 0.59. This deviation is likely to be due to the underlying high variance of gene expression data generated by RACIPE. Nonetheless, the EMT metric has shown a consistent characterization results of the RACIPE-generated samples for the hybrid E/M I, II and M categories. We also find that the mean hybrid E/M I signature is close to that of epithelial samples, while the mean hybrid E/M II signature localizes closer to that of mesenchymal samples in the predictor space (**Fig. 2b**), which is consistent with the clustering results observed in **Fig. 1b**. Interestingly, the hybrid E/M II samples have similar mean VIM/CDH1 levels as that of epithelial samples and similar mean CLDN7 levels as that of mesenchymal samples, thus sharing features of both epithelial and mesenchymal signatures (Fig. 2b). This observation is reminiscent of a recent study showing that overexpression of miR-200 in mesenchymal subpopulation of HMLE cells induced only a partial EMT - the mesenchymal score of these cells did not change upon miR-200 overexpression, but the epithelial score increased [38].

**Figure 2.**
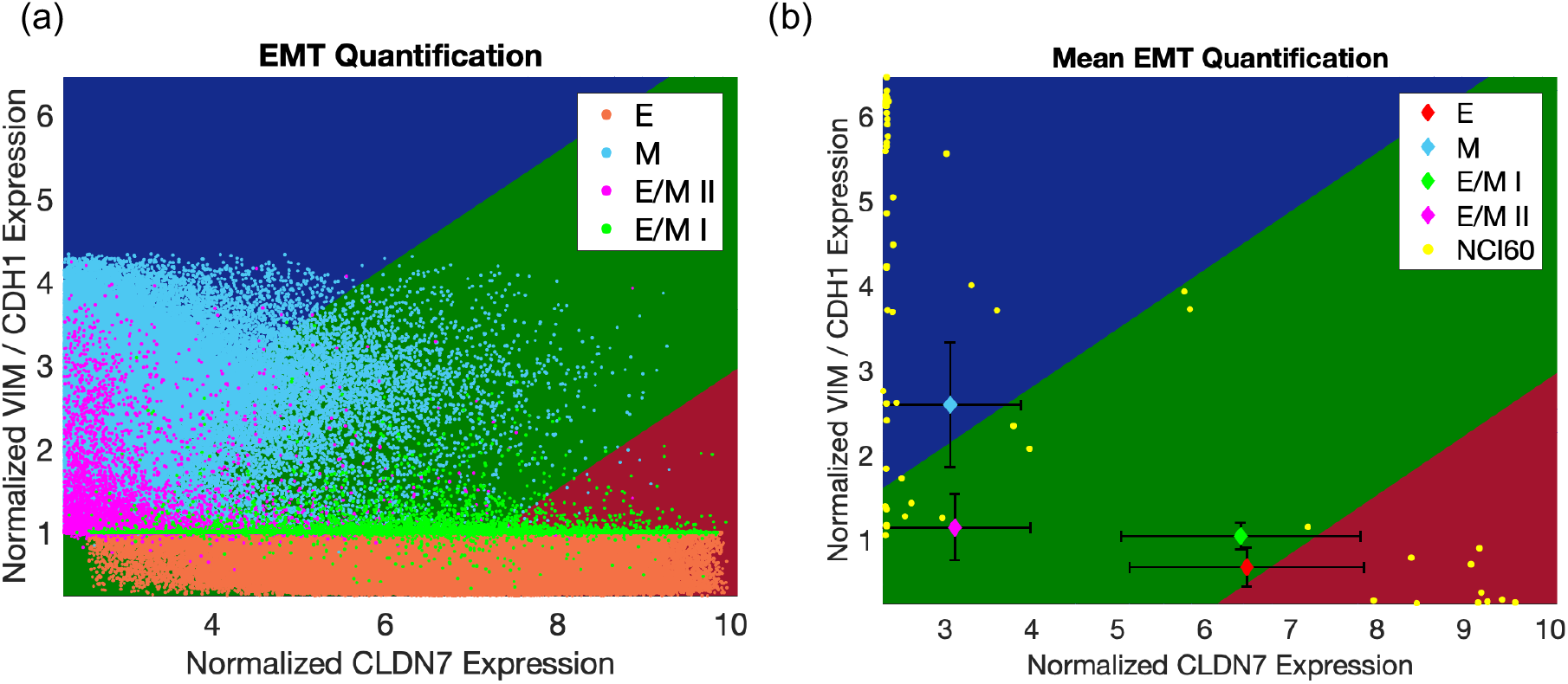
**Characterization of the RACIPE-generated samples by the EMT metric**. (a) Population level projection of RACIPE generated data onto EMT metric predictor space. VIM, CDH1, and CLDN7 expression values for each sample generated from RACIPE are used to plot various populations categorized by RACIPE (red region of EMT metric predictor space corresponding to E, dark green corresponding to hybrid E/M I and hybrid E/M II, and blue corresponding to M; salmon identifying RACIPE-generated E samples, light green identifying hybrid E/M I samples, magenta identifying hybrid E/M II samples and sky blue identifying M samples). (b) Predicted mean expression signature for each RACIPE population cluster. Each colored diamond represents the mean expression of RACIPE-predicted hybrid E/M I (light green), hybrid E/M II (magenta), E (salmon), and M (sky blue) samples, plotted alongside the NCI-60 training dataset (yellow). Black error bars represent standard deviations along each predictor dimension.

### Using the EMT scoring metric to distinguish between samples comprised of individual hybrid E/M cells and samples that are mixtures of E and M cells

After establishing this initial success in predicting the population phenotype, we evaluated the ability of the EMT scoring metric to distinguish between populations consisting mostly of individual hybrid E/M cells versus samples that are mixtures of E and M cells. Unmixed samples of epithelial (EMT score 0 ≤ *μ* < 0.5) or unmixed mesenchymal (EMT score 1.5 < *μ ≤* 2) cells are easily characterized by the EMT metric, which localizes their positions on the EMT predictor space to the red and blue regions, respectively **(Fig. 3)**. Categorizing a hybrid E/M (EMT score 0.5 ≤ *μ ≤* 1.5) sample is less straightforward as there are multiple possibilities that explain such a prediction. The sample may: a) contain individual hybrid E/M cells, b) contain a mixture of epithelial and pure mesenchymal cells, or c) some combination of epithelial, mesenchymal, and hybrid E/M cells. As shown in **Fig. 4**, mixtures of RACIPE-generated epithelial and mesenchymal samples may be well-approximated by a convex combination curve in the two-dimensional predictor space, hereafter referred to as a ‘mixture curve’, connecting the positions of mean signatures of pure epithelial and pure mesenchymal RACIPE-generated samples. Interestingly, the position of mean RACIPE-generated hybrid E/M signatures, particularly E/M II samples, concentrates further away from the mixture curve. Moreover, the higher percentage of the individual hybrid E/M II cells in the sample, the further the position of the sample from the mixture curve in the predictor space (Fig. 3). These results indicate that the EMT scoring metric may be used to predict the proportions of different subpopulations of samples that are mixtures of different EMT phenotypes.

**Figure 3.**
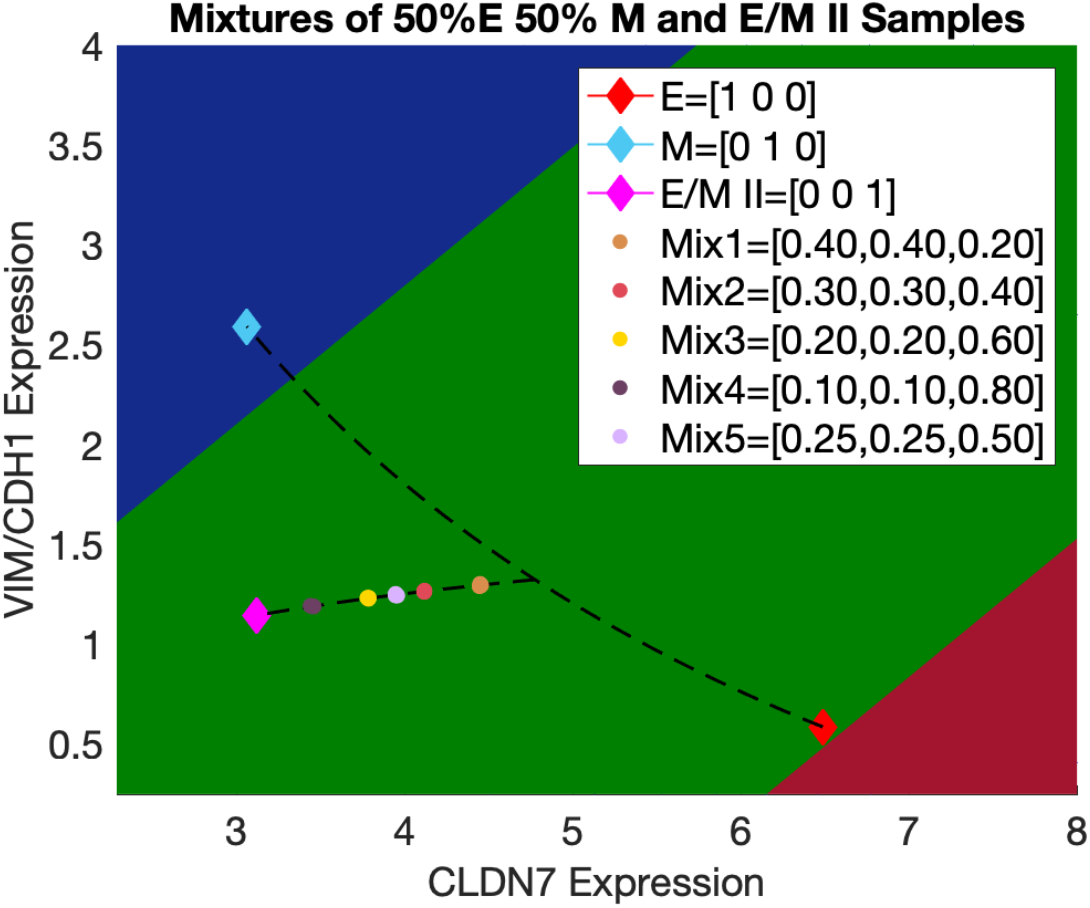
**Use of the EMT metric to characterize the mixtures of E, M and hybrid E/M samples where the proportion of E is equal to that of M in each mixture**. Predicted mixture proportions contained in each mixture {E, M, E/M II} are reported in the figure legends for each sample and predicted based on their projection to the convex mixture curve adjoining E/M II and the midpoint of E and M.

### The EMT scoring metric can accurately resolve mixtures of pairs of EMT phenotypes

As the EMT metric has shown its power in distinguishing between individual hybrid E/M cells and mixtures of E and M cells, we evaluated the ability of the EMT scoring metric to resolve the proportions of two different EMT phenotypes mixed in varying ratios. Using RACIPE, we generated three pairs of *in silico* mixtures – a) mixtures of E and M samples, referred to as {E, M}, b) mixtures of E and hybrid E/M II samples, referred to as {E, E/M II} and c) mixtures of M and hybrid E/M II samples, referred to as {M, EM II} with five different ratios (see **Materials and Methods** for details). We used hybrid E/M II for generating mixtures because the mean expression signature of hybrid E/M II exhibits similar CLDN7 expression as that of a M phenotype and similar VIM/CDH1 level as that of an E phenotype, and is thus more likely to be a canonical representative ‘hybrid’ E/M phenotype. For every fixed ratio of a given pair of phenotypes, three independent replicates were generated, and they showed highly reproducible behavior (**Fig. 4**). The EMT metric-estimated proportions of E and M in {E, M} are [49%, 51%], [40%, 60%], [59%, 41%], [20%, 80%], [79%, 21%] with respective mean absolute errors of [0.44%, 0.45%, 0.55%, 0.86%, 1.37%] across replicates. These estimates are very close to the true proportions of E and M that are [50%, 50%], [40%, 60%], [60%, 40%], [20%, 80%], [80%, 20%], respectively (**Fig. 4a, d**). Similarly, the EMT metric-estimated proportions of E and hybrid E/M II in {E, E/M II} are converge to their true values of [50%, 50%], [60%, 40%], [40%, 60%], [20%, 80%], [80%, 20%] with respective mean errors of [0.26%, 0.25%, 0.23%, 0.04%, 0.11%] (Fig. 3b, d). Finally, the EMT metric-estimated proportions of M and hybrid E/M II in {M, E/M II} are [50%, 40%], [60%, 40%], [40%, 60%], [20%, 80%], [80%, 20%] across replicates with respective mean errors of [0.26%, 0.08%, 0.18%, 0.11%, 0.51%], again converging to their true values (Fig. 3c, d). Together, these results indicate that the EMT metric can effectively deconvolve mixtures of multiple EMT phenotypes, and thereby accurately estimate their proportions in a given mixture containing two different EMT phenotypes.

**Figure 4.**
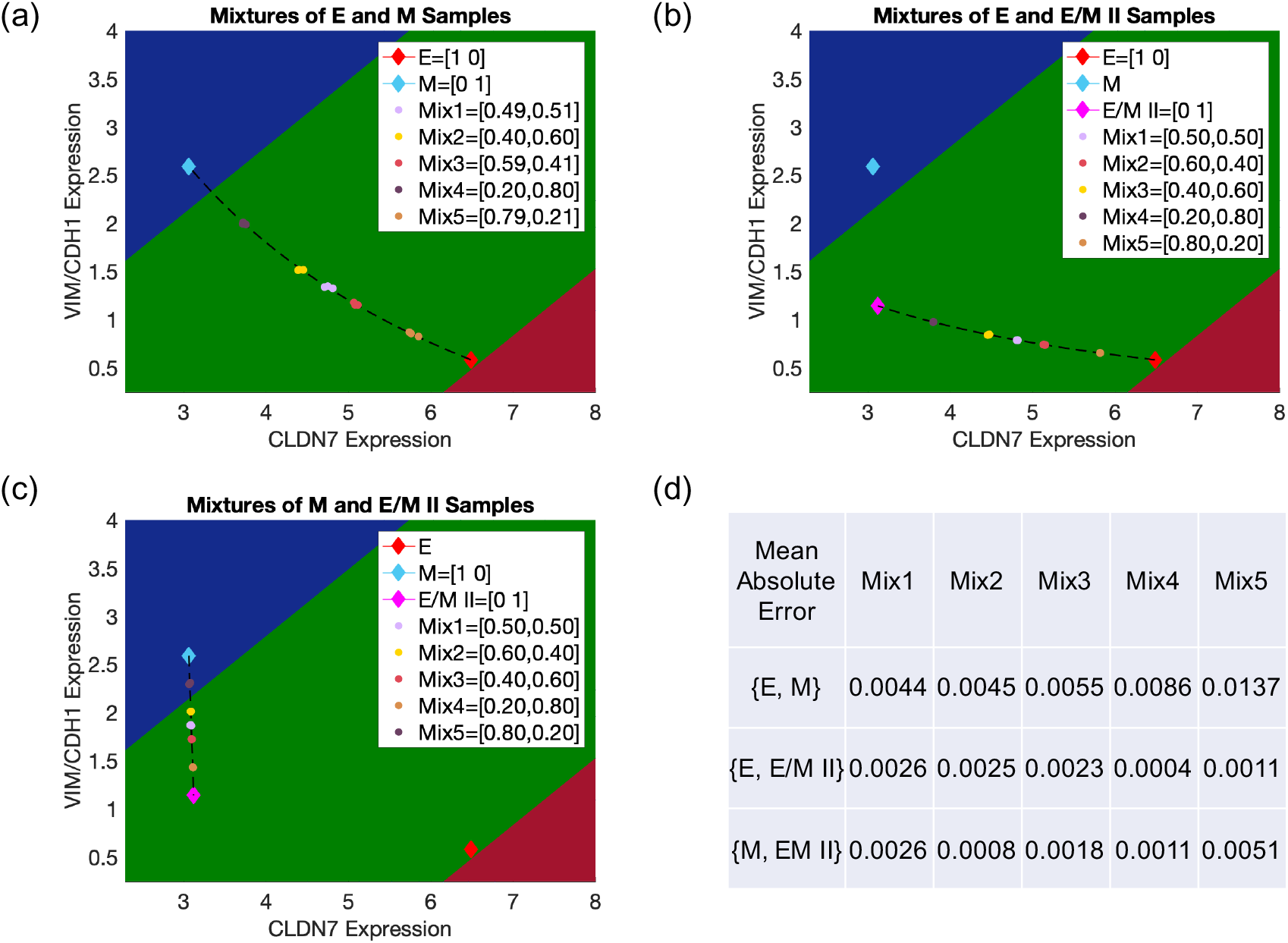
**Use of the EMT scoring metric to characterize the mixtures of two different EMT phenotypes**. Predicted mixture proportions contained in each mixture {E, M} (a), {E, E/M II} (b) and {M, E/M II} (c) are reported in the figure legends for each sample and predicted based on their projection to the curve of convex combinations for each non-mixed sample pair. In all cases, all three replicates for one mixture are tightly distributed and therefore have significant overlap. (d) The mean absolute error of predicted mixture proportions. The true mixture proportions of {E, M} are [50%, 50%] (Mix1) [40%, 60%] (Mix2) [60%, 40%] (Mix3) [20%, 80%] (Mix4) and [80%, 20%] (Mix5). The true mixture proportions of {E, E/M II} and {M, E/M II} are [50%, 50%] (Mix1) [60%, 40%] (Mix2) [40%, 60%] (Mix3) [20%, 80%] (Mix4) and [80%, 20%] (Mix5).

Motivated by the initial observation that the mean RACIPE-generated epithelial signature was binned as hybrid E/M, we further analyzed whether a mixture model constructed on samples binned as epithelial by the EMT metric would lead to significantly different results. We performed sample purification and isolated the purified epithelial samples that exhibit high expression of multiple E markers including CDH1, GRHL2, OVOL2, ΔNp63α and CLDN7. All of these players are known to be key regulators of epithelial phenotype [39–42]. Similarly, we performed sample purification to isolate the purified mesenchymal samples that have high expression of multiple M markers, including VIM, ZEB1, ZEB2, TGF-β and FOXC2. These markers are experimentally known as key inducers of the mesenchymal phenotype [26,43]. This step is the *in silico* analogue to flow cytometric sorting of E (and M) samples for a more robust characterization of these two cell states, as captured by the increased accuracy of EMT metric in categorizing these samples (**Tables S2, S3**). The purified epithelial and mesenchymal samples are accurately predicted via EMT metric (*P_E_* = 0.82, *μ* = 0.18 and *P_M_* = 0.88, *μ* = 1.88, respectively), and visually fall well within their respective domains of predictor space (**Fig. S1, See SI sections 2-3 for more details**). Mixtures containing purified E and M samples for five different ratios were synthesized and analyzed by the EMT metric. Similar to the case with ‘unpurified’ samples (i.e. using all E samples), the EMT metric accurately predicts the proportions of the mixtures of ‘purified’ E and M samples (**Fig. S2**). In short, the sample purification analysis here was only to offer alternative reference points for E and M signatures which might be more representative of real biological samples.

### The EMT scoring metric can accurately resolve mixtures of hybrid E/M, E and M phenotypes when E and M phenotypes have the same proportions

As the first step to evaluate the ability of the EMT metric in resolving mixtures of E, hybrid E/M and M phenotypes, we use the RACIPE-generated E, hybrid E/M II and M gene expression data to synthesize mixtures of all three phenotypes, referred to as {E, M, E/M II}. We keep the proportions of E and M samples the same and vary the proportions of the hybrid E/M II samples when preparing {E, M, E/M II}. Mixtures with five different ratios were randomly selected in triplicate and were highly reproducible and resolvable (Fig. 4). The EMT metric-estimated proportions of E, M and E/M II in {E, M, EM II} are [25.5%, 25.5%, 49%], [30%, 30%, 40%], [20%, 20%, 60%], [10%, 10%, 80%], [40%, 40%, 20%] with respective mean errors of [0.32%, 0.2%, 0.19%, 0.37%, 0.22%] across replicates which are very close to the true proportions of E, M and E/M II: [25%, 25%, 50%], [30%, 30%, 40%], [20%, 20%, 60%], [10%, 10%, 80%], [40%, 40%, 20%], respectively (**Fig. 3**). These results lead to the hypothesis that samples dominated by the hybrid E/M II phenotype tend to localize far from the ‘mixture curve’, while samples locating close to the ‘mixture curve’ are possible mixtures of E and M cells.

### Using the EMT scoring metric to characterize the EMT proportions of NSCLC cell lines

To test the hypothesis that samples dominated by the hybrid E/M phenotype localize far from the ‘mixture curve’, we performed experiments in multiple NSCLC cell lines that have been categorized as epithelial, mesenchymal, or hybrid E/M based on bulk proteomic measurements (GSE63882) [36]. The EMT scoring metric characterizes H1568 cell line as largely epithelial, H1792 cell line as largely mesenchymal, and the H920, H1944, H2228 and HCC2279 cell lines as hybrid E/M (**Fig. 5a**). Among the hybrid E/M cell lines, H920 and H2228 locate far from the ‘mixture curve’ in the EMT metric predictor space, thus being predicted to contain a large proportion of the hybrid E/M cells, while H1944 and HCC2279 cell lines lie on/close to the curve, which means that they are possible mixtures of E and M cells.

To validate this prediction, we conduct single-cell immunofluorescence experiments to analyze the presence of canonical epithelial and mesenchymal markers - E-cadherin and vimentin - in these cell lines. H1568 cells largely stain for E-cadherin, with only a few cells staining for vimentin, thus depicting a predominantly epithelial phenotype. Conversely, H1792 cells express only vimentin, and very low E-cadherin, exhibiting a predominantly mesenchymal phenotype (**Fig. 5b**). Most cells in H920 and H2228 tend to co-express E-cadherin and vimentin and some cells express only E-cadherin, thus demonstrating a mixture of hybrid E/M and epithelial cells, with a hybrid E/M phenotype being the dominant subpopulation (**Fig. 5b**). The H1944 cell line contains cells expressing either only E-cadherin or only vimentin thus being a mixture of E and M cells. This is consistent with the above EMT score finding. On the other hand, most individual cells in HCC2279 cell line co-express E-cadherin and vimentin, thus being hybrid E/M cells. Results of H1944 and HCC2279 suggest that a cell line lying on/close to the mixture curve might be a mixture of E and M but also might be a single-cell hybrid that just happens to lie on that line. Put together, this *in vitro* characterization of these six NSCLC cell lines offer a first proof-of-principle validation of our predictions about their EMT-ness, estimated by their relative distance to the ‘mixture curve’ in the predictor space of the EMT scoring metric.

**Figure 5.**
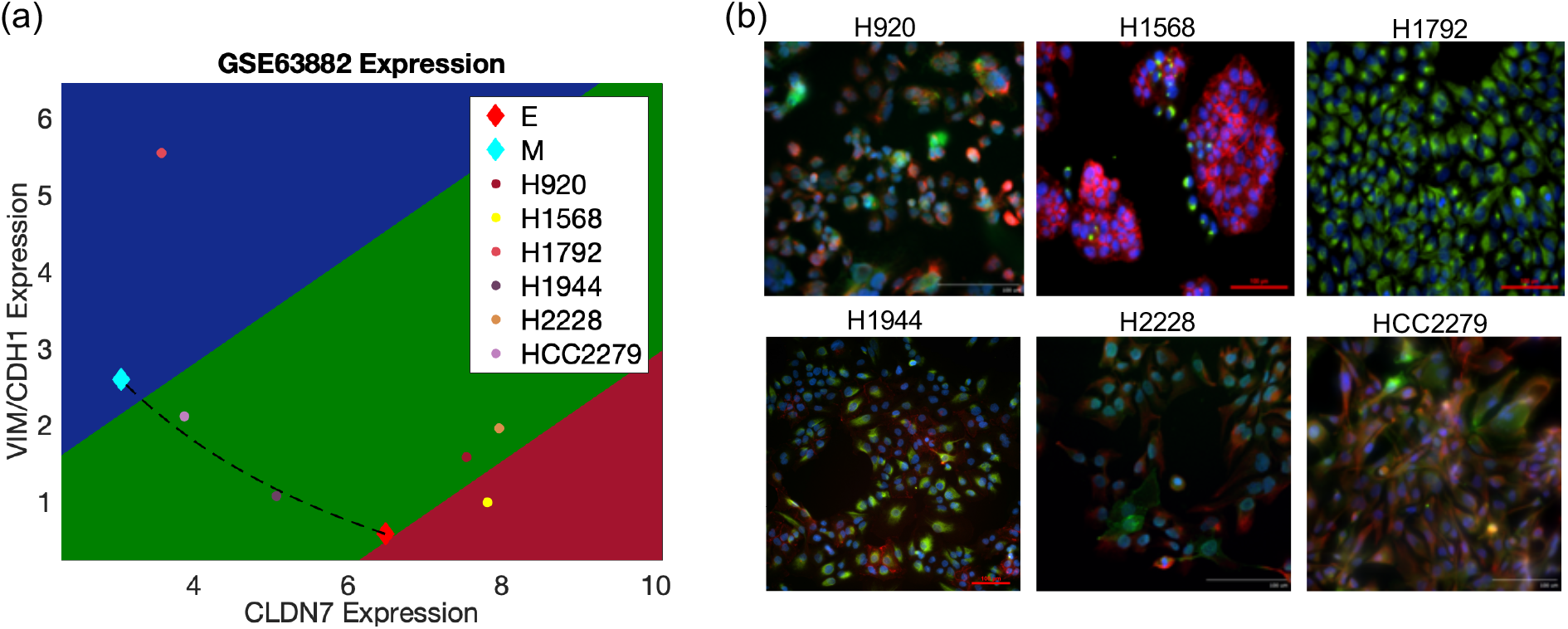
**Characterizing the EMT phenotypes. (**a) EMT expression signatures for H920, H1568, H1792, H1944, H2228, and HCC2279 are projected into EMT predictor space relative to the mixture curve between RACIPE-generated E and M samples. (b) Expression of E-cadherin (red) and vimentin (green) examined by immunofluorescence staining. Scale bar 100 μm.

## Discussion

The existence and characterization of one or more hybrid epithelial/mesenchymal (E/M) phenotypes accessible during the epithelial-mesenchymal transition has generated recent attention due to their association with high metastatic potential and cancer stemness [35,44–46]. Increasing evidence suggests that cancer cells may acquire a spectrum of hybrid E/M phenotypes, characterized by varying extents of epithelial and mesenchymal traits [35]. Thus, a rigorous characterization of the EMT status of tumors, especially the probability that a given sample is hybrid E/M-like, may contain significant prognostic value and may hold avenues for better targeted therapeutic developments [47].

Recent years have seen a surge in various computational and experimental approaches to quantitatively characterize various aspects of EMT, including characterization of hybrid E/M phenotypes [17–21,25,26,27–29,35,36,44,45]. Here, we test our EMT scoring metric by applying it to the *in silico* gene expression data generated by RACIPE. We show that the hybrid E/M II phenotype is effectively distinguished from mixtures of E and M samples, based on their different localizations in the predictor space generated by the EMT scoring metric. As compared to another EMT scoring method that requires data of the entire transcriptome [48], our approach presented here can quantify a sample‘s ‘EMT-ness’ using a very small set of predictors with reasonable categorization results.

Our results presented here may suffer from multiple limitations. First, the EMT regulatory network constructed here, although based on extensive literature search, is in no way complete. For the purpose of computational cost, we focused on a conserved network implicated in EMT in multiple contexts. However, there may be crucial connections that we have inadvertently missed. Second, we simulated both transcriptional factors and microRNA-dependent mechanisms via the shifted Hill function formulation, while their detailed dynamics may be quite different [25]. That means we must determine the EMT status of RACIPE models only by the “expression” data as opposed to using protein concentration and/or actual cell shape and behaviors. It is worth noting that EMT is a multi-dimensional process including changes in not only gene expression, but also in many biophysical properties such as cell polarity [49]. Thus, future studies should identify biophysical features that, when combined with gene expression patterns, might provide further refinement in resolving hybrid E/M phenotypes. Third, the training dataset used for EMT scoring metric are bulk-cell gene expression profiles of 59 cancer cell lines, and 37 out of the 59 are mesenchymal [50]. Further detailed experimental characterization of the EMT phenotype at a single-cell level [51] can help expand the training dataset and perhaps increase the sensitivity, specificity and accuracy in quantifying the EMT spectrum. Despite these weaknesses, we feel that we have succeeded in showing broad consistency between the RACIPE and the EMT scoring metric and also generated a test for the existence of single cell hybrids.

The ability to fully distinguish between mixtures of E and M cells and purely hybrid E/M samples remains a relevant challenge as both of these share a common hybrid gene expression pattern on the population level. By projecting the mixtures and purely hybrid E/M cells into the predictor space of the EMT metric, we find that the mixtures of epithelial and mesenchymal tend to cluster along the convex curve adjoining purely epithelial and purely mesenchymal while samples containing purely hybrid E/M cells are not necessarily restricted to the mixture curve between unmixed epithelial and mesenchymal samples, as shown in **Figs. 3** and **4**. Based on this observation, we speculate that hybrid E/M cells and mixtures of epithelial and mesenchymal cells may often be distinguishable. Gene expression data of more cell lines with a well-characterized EMT status, i.e. purely epithelial, mesenchymal, hybrid E/M or mixtures of well-known proportions of these phenotypes, can be used to test this hypothesis.

## Acknowledgements

This work is supported by National Science Foundation (NSF) grants - Center for Theoretical Biological Physics NSF PHY-1427654 and PHY-1605817. J.T. George is also supported by the National Cancer Institute of the National Institutes of Health F30CA213878. M.K. Jolly is also supported by a training fellowship from the Gulf Coast Consortia on the Computational Cancer Biology Training Program CPRIT Grant No. RP170593.

